# Altricial, but not unusual: Human developmental timing follows general mammalian life-history scaling

**DOI:** 10.64898/2026.06.16.732383

**Authors:** Can Dora Akcan, Doğa Keçe, Kaan Kerman

## Abstract

Humans are widely regarded as unusually slow to develop, exhibiting prolonged childhood and extended dependence on caregivers. However, this view is based primarily on comparisons with other primates, leaving unresolved whether humans remain distinctive within the broader diversity of mammals. We addressed this question by situating human development in a comparative framework using gestation length, weaning age, and age at sexual maturity for both sexes across 462 mammalian species representing 25 orders. Each trait was examined both as an absolute value and as a proportion of the longest verified captive lifespan. In absolute terms, human developmental traits fell within the upper range of mammalian variation. When expressed relative to lifespan, however, gestation shifted toward the lower end of the distribution, whereas weaning age and sexual maturity occupied intermediate positions, indicating that human developmental timing largely follows general mammalian scaling patterns rather than representing a pronounced outlier. These findings suggest that key features of human dependency are better understood as extensions of broader evolutionary trends than as uniquely human life-history characteristics.

## 1. Introduction

Life histories across primates, particularly among great apes, are characterized by relatively slow developmental schedules and extended lifespans compared to most mammals (Kappeler & Pereira, 2003; van Schaik et al., 2006). Within this context, humans are frequently described as unusually altricial (Portmann, 1969; Dunsworth et al., 2012). A key feature of this interpretation is an extended postnatal developmental period in humans, which results in prolonged dependency on caregivers for both survival needs and the development of sociocognitive skills relevant to human eusociality and cooperative care (Bogin, 1999, 2001; Hrdy, 2009; Kaplan et al., 2000; Bogin & Varea, 2017). A similar pattern has also been observed in studies focusing solely on the primate order, where humans exhibit distinctive life history parameters such as reduced adult mortality, shorter interbirth intervals, large neonatal body mass, extended juvenile dependency, earlier weaning, and prolonged post-reproductive longevity (Robson et al., 2006; Robson & Wood, 2008; DeSilva, 2011; Miller et al., 2019).

Nevertheless, the developmental characterization of humans as potentially atypical in terms of dependency may require further investigation. Human patterns largely follow and, in some respects, extend this pattern (Charnov & Berrigan, 1993). Also, while humans are frequently described as maturing late, comparative analyses suggest that age at first reproduction is consistent with great ape expectations once body size and phylogenetic relationships are taken into account (Robson et al., 2006; Miller et al., 2019). Indeed, recent evidence indicating later maturation in chimpanzees than previously estimated reduces the apparent distinctiveness of human developmental timing (Walker et al., 2018; Miller et al., 2019). Moreover, while human infants are born with relatively small brains due to birth constraints from bipedalism (Gómez-Robles et al., 2023), the degree of altriciality appears similar in humans and apes (Schultz, 1961; Charvet et al., 2023). Even “emerging adulthood,” often described as uniquely human (Hochberg & Konner, 2020), parallels stages seen in other primates (Goodall, 1986).

Beyond primates, life history variation across mammals spans a well-documented fast-slow continuum, ranging from rapid developers with short lifespans to slow-maturing, long-lived species (Promislow & Harvey, 1990; Read & Harvey, 1989). Several non-primate taxa exhibit unexpectedly slow life histories given their body size or phylogenetic position. These include bats, whose maximum recorded lifespans are on average 3.5 times greater than those of similarly sized non-flying placentals (Austad & Fischer, 1991; Wilkinson & South, 2002); hibernating mammals, which show longer lifespans, later maturation, and slower reproductive rates than non-hibernating counterparts of equivalent size (Turbill et al., 2011); and elephants, which exhibit late reproductive onset and extended post-reproductive survival (Lee et al., 2016; Lahdenperä et al., 2014). Despite these cases, systematic comparative investigations that situate humans within this broader mammalian framework remain relatively limited. Among odontocetes, post-reproductive lifespans have been documented in killer whales and short-finned pilot whales (Foote, 2008), with subsequent work extending this to belugas and narwhals as well (Ellis et al., 2018).

Crucially, most studies placing humans in a comparative life history context have focused on primate-only samples (Robson et al., 2006; Robson & Wood, 2008; Miller et al., 2019), leaving open the question of whether human developmental timing appears distinctive when evaluated against the full range of mammalian variation rather than against close phylogenetic relatives alone. Without such a comparison, it is difficult to determine whether humans represent a genuine outlier in mammalian life history space, or whether they fall within the broader continuum of slow strategies distributed across long-lived taxa.

In the present study, we wanted to assess the extent to which humans can be characterized as highly altricial by situating human developmental traits within a comparative mammalian framework. To that end, we examined gestation length, weaning age, and age at sexual maturity (for both sexes) as proportions of maximum lifespan across a large sample of mammalian species (N = 462) spanning 25 orders, thereby enabling standardized comparisons across taxa with widely differing life histories. We applied a descriptive analytical framework to the dataset to evaluate whether human developmental timing is consistent with expected variation or appears as an outlier within the broader continuum of mammalian life history strategies. We used both absolute and proportional values in the analysis.

## 2. Methods

### 2.1. Taxonomic scope and data sources

We compiled the life history data across 462 non-human mammalian species based on the recent taxonomic classification. We used peer-reviewed sources to obtain reliable documentation of life history metrics for each species, and omitted species from the list when the sources were unavailable, incomplete, or inconsistent (see section 2.2.). For humans, we relied on the most up-to-date sources on the topic (Peters & Kemper, 2012; Dettwyler, 2004; Jukic et al., 2013).

Two of the authors (D.K. and C.D.A.) performed the task of data extraction independently, each compiling values into separate spreadsheets prior to cross-comparison. We identified discrepancies between extractions through systematic comparisons and resolved by joint review of the original source text. Where the source remained ambiguous following review, we excluded the relevant species–trait combination from all analyses rather than resolved by averaging or imputation.

### 2.2. Inclusion and exclusion criteria

We applied the inclusion and exclusion criteria independently to each species-trait combination (Figure 1). This allowed us to retain valid data for some variables while excluding the same species for other missing variables, hence maximizing the usable dataset without introducing systematic bias through listwise exclusion.

**Figure 1.**
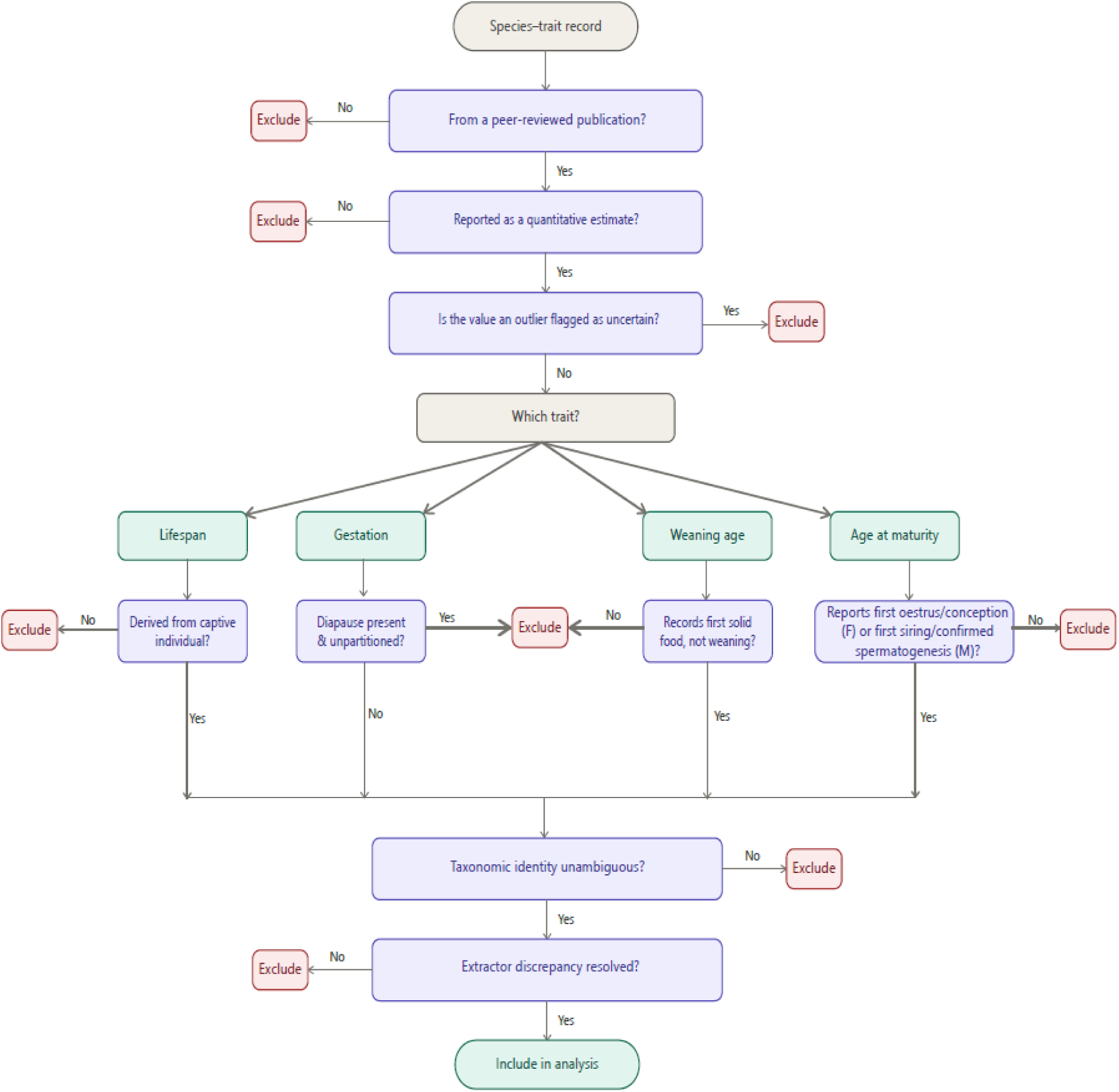
Inclusion-exclusion decision tree. Nodes represent criteria applied sequentially to each species–trait combination.

We included a record of life history variable only if all the following conditions were satisfied:

- Life-history data is published in a peer-reviewed journal.
- Life history data is a quantitative estimate (i.e., a single measurement from an individual or a measure of central tendency, including a mean or a median).
- In the context of lifespan, data is derived from captive individuals or populations, where external sources of mortality are largely controlled or minimized (e.g., predation, resource limitation, parasite load).

We excluded data if any of the following conditions applied:

- Life history data is derived from a source that is not published in a peer-reviewed paper and cannot be verified against an independent source (e.g., a studbook entry without an associated publication, a zoo record without documentation of methodology)
- Data is flagged as uncertain or exceptional by the source author.
- For gestation length, the reported duration includes the period of embryonic diapause (delayed implantation) and no partitioned estimate of active gestation is available.
- For weaning age, the record described age at first supplemental solid food intake rather than age at complete cessation of nursing, and no estimate of the latter was available in the same or a corroborating source.
- The taxonomic identity of the study animal is ambiguous or was contested at the time of this study.

### 2.3. Life history & dependency variables

We operationalized three life history traits for the purposes of this study.

*Gestation length,* defined as the mean duration of active intrauterine development from implantation to parturition, measured in days (Jones et al., 2009; Renfree & Shaw, 2000). For species exhibiting embryonic diapause, we retained only the period of active gestation following implantation when such partitioned data were available.

*Weaning age,* defined as the age, in days post-partum, at which offspring achieve complete nutritional independence from maternal milk (Jones et al., 2009; Lee, 1996). Where sources reported a range without a central estimate, the mean was used. For species in which a period of partial nursing alongside solid food intake was documented, we used the age at complete cessation of suckling in preference to the age at first supplemental feeding.

We represented *age at reproductive maturity* for each sex separately. For females, we defined it as the age at first estrus or first successful conception. For males, we defined it as age at first successful copulation resulting in offspring, or, where behavioral reproductive data were unavailable, as the age at onset of spermatogenesis confirmed by histological examination. Where a source reported ‘sexual maturity’ without specifying sex, we assigned the same value to both sexes and flagged it in the database; such records were retained only where they were consistent with sex-specific values from independent sources for the same species.

All temporal variables were converted to days prior to analysis. To facilitate cross-species comparisons that are independent of absolute lifespan magnitude, we also computed proportional indices by dividing each trait value by the species’ maximum captive lifespan, defined as the longest verified survival in a captive setting. We accepted captive lifespan records described as estimates or inferred from dental or skeletal ageing methods only when the estimation method was explicitly stated in the source material.

### 2.4. Statistical analysis

We computed descriptive statistics for each developmental variable both in terms of its absolute value and in relation to its proportional effect, followed up by documenting the percentile ranks for the human data point within the distribution of each measured trait. To facilitate visual comparison, we log-transformed count/frequency data (i.e., age) and logit-transformed proportional data (dependency in relation to maximum lifespan). We implemented all statistical analyses in Python (v3.8.10; Python Software Foundation), using the pandas (v2.0.3), NumPy (v1.24.4), and matplotlib (v3.7.5) libraries.

## 3. Results

### 3.1. Analyzed Mammalian Taxa

Implementing the first phase of inclusion-exclusion criteria for detecting at least one credible reliable dependency trait resulted in 462 species across 25 orders. Most of the taxa belonged to the order Rodentia, followed by Artiodactyla and Primates.

Following application of all inclusion and exclusion criteria, the final count varied in relation to the life history trait. Gestation length was the most frequently recorded developmental metric in the literature (n = 304) followed by the age of meaning (n = 254). Age at reproductive maturity was reported more commonly in females (n = 256) than males (n = 221), likely reflecting measurement bias. The complete list of included species, with all retained trait values and their original sources, can be accessed in the supplementary material (Datasets S1 and S2).

### 3.2. Dependency Metric Distributions

Descriptive statistics for all dependency distributions are presented in Table 1.

**Table 1.**
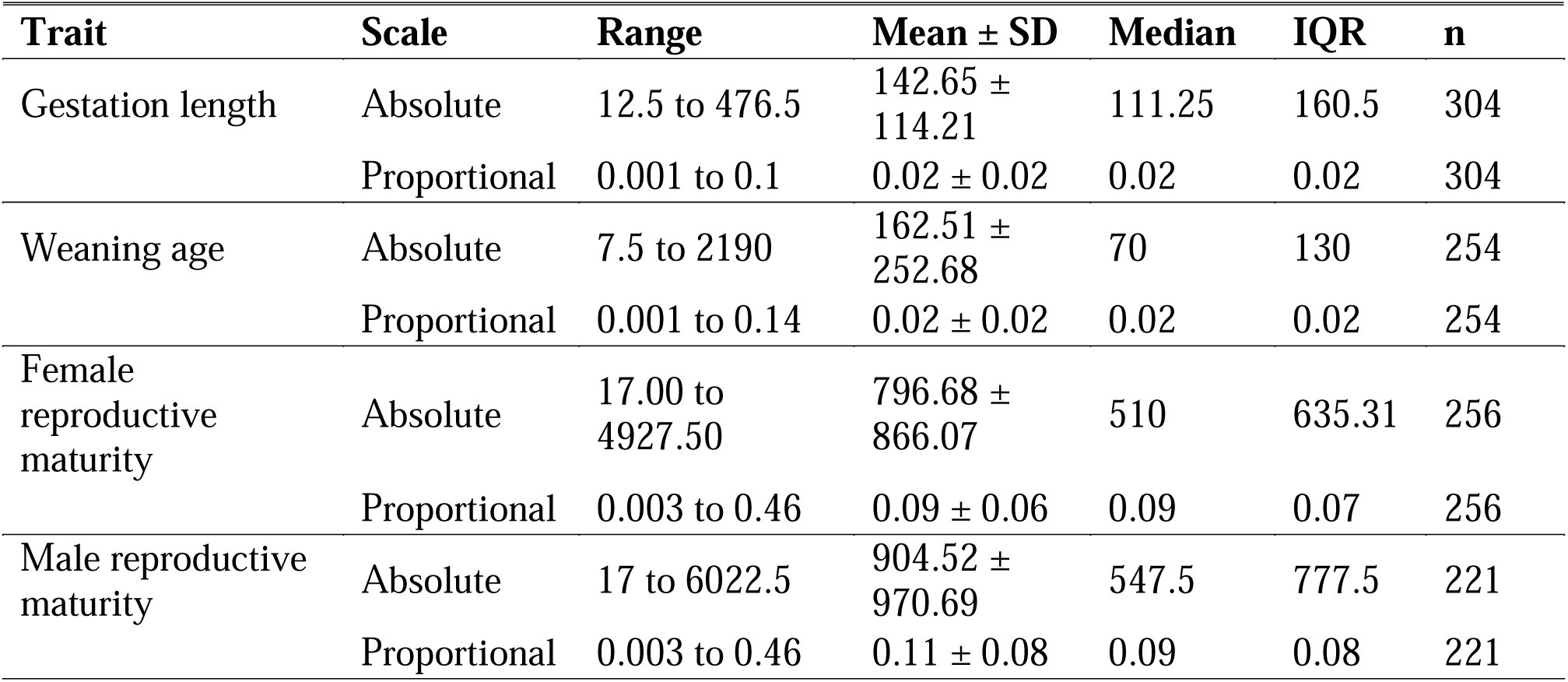
Summary statistics for mammalian life-history traits included in the analyses. For each trait, descriptive statistics are reported for absolute values and lifespan-standardized proportional values. Values shown are range mean ± standard deviation (SD), median, interquartile range (IQR), and sample size (n).

| Trait | Scale | Range | Mean ± SD | Median | IQR | n |
| --- | --- | --- | --- | --- | --- | --- |
| Gestation length | Absolute | 12.5 to 476.5 | 142.65 ± 114.21 | 111.25 | 160.5 | 304 |
| | Proportional | 0.001 to 0.1 | 0.02 ± 0.02 | 0.02 | 0.02 | 304 |
| Weaning age | Absolute | 7.5 to 2190 | 162.51 ± 252.68 | 70 | 130 | 254 |
| | Proportional | 0.001 to 0.14 | 0.02 ± 0.02 | 0.02 | 0.02 | 254 |
| Female reproductive maturity | Absolute | 17.00 to 4927.50 | 796.68 ± 866.07 | 510 | 635.31 | 256 |
| | Proportional | 0.003 to 0.46 | 0.09 ± 0.06 | 0.09 | 0.07 | 256 |
| Male reproductive maturity | Absolute | 17 to 6022.5 | 904.52 ± 970.69 | 547.5 | 777.5 | 221 |
| | Proportional | 0.003 to 0.46 | 0.11 ± 0.08 | 0.09 | 0.08 | 221 |

All three life history traits exhibited positive skewness in both their absolute values and their proportions relative to maximum lifespan (Figure 2). In terms of absolute metrics, human dependency measurements fell within the fourth quartile of the overall mammalian distribution: Human gestation length (268 days) corresponded to the 86.18th percentile, weaning age (912.50 days) to the 98th percentile, male age at reproductive maturity (4161 days) to the 98.6th percentile, and female age at reproductive maturity (4927.50 days) to the 99.8th percentile. On the proportional scale, human dependency metrics were substantially lower relative to their positions in the absolute distribution. Gestation length fell at the 4.6th percentile (proportion = 0.01), weaning age at the 64.9th percentile (proportion = 0.02), and age at reproductive maturity at the 71.1st percentile for females (proportion = 0.11) and the 52.5th percentile for males (proportion = 0.09).

**Figure 2.**
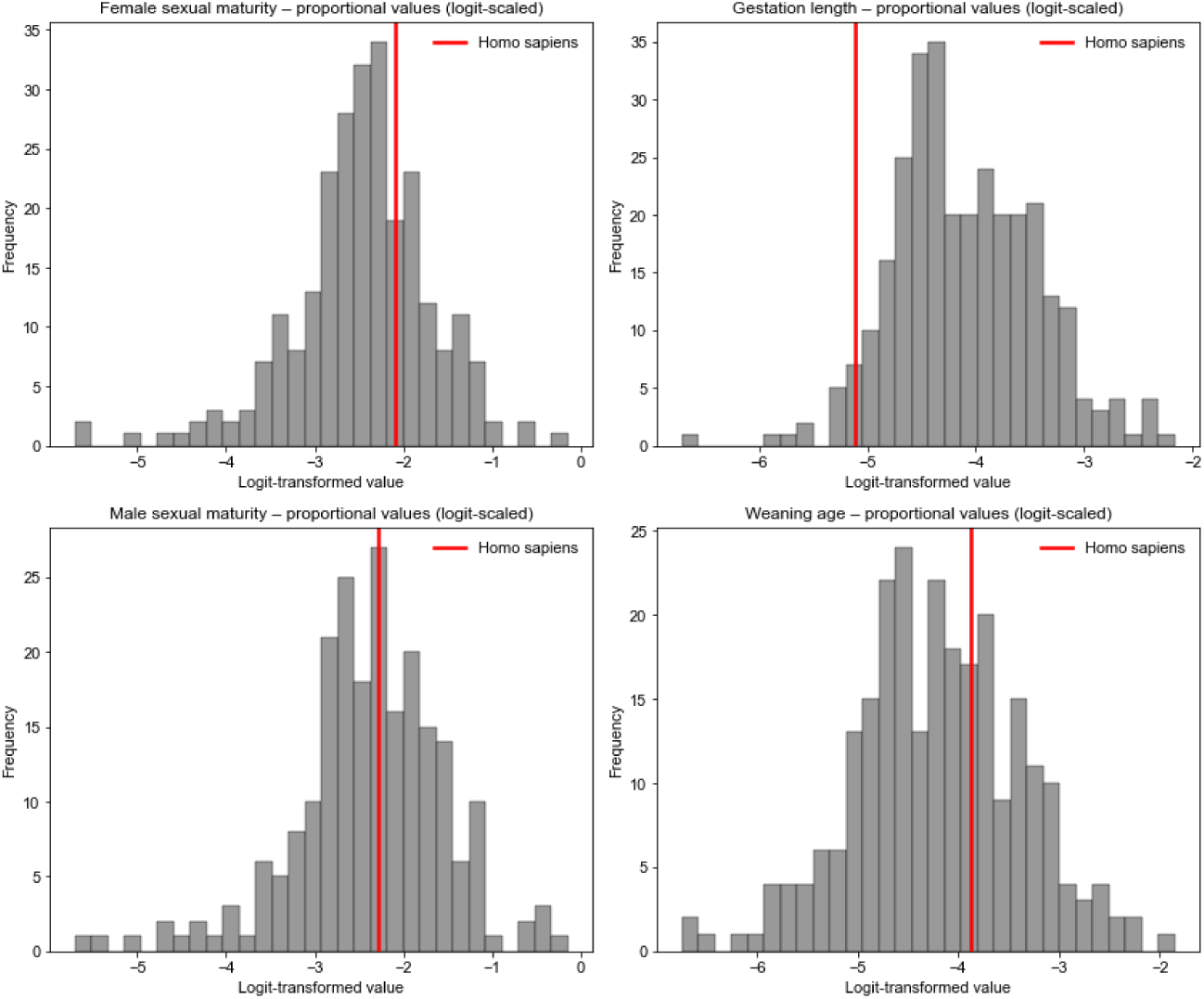
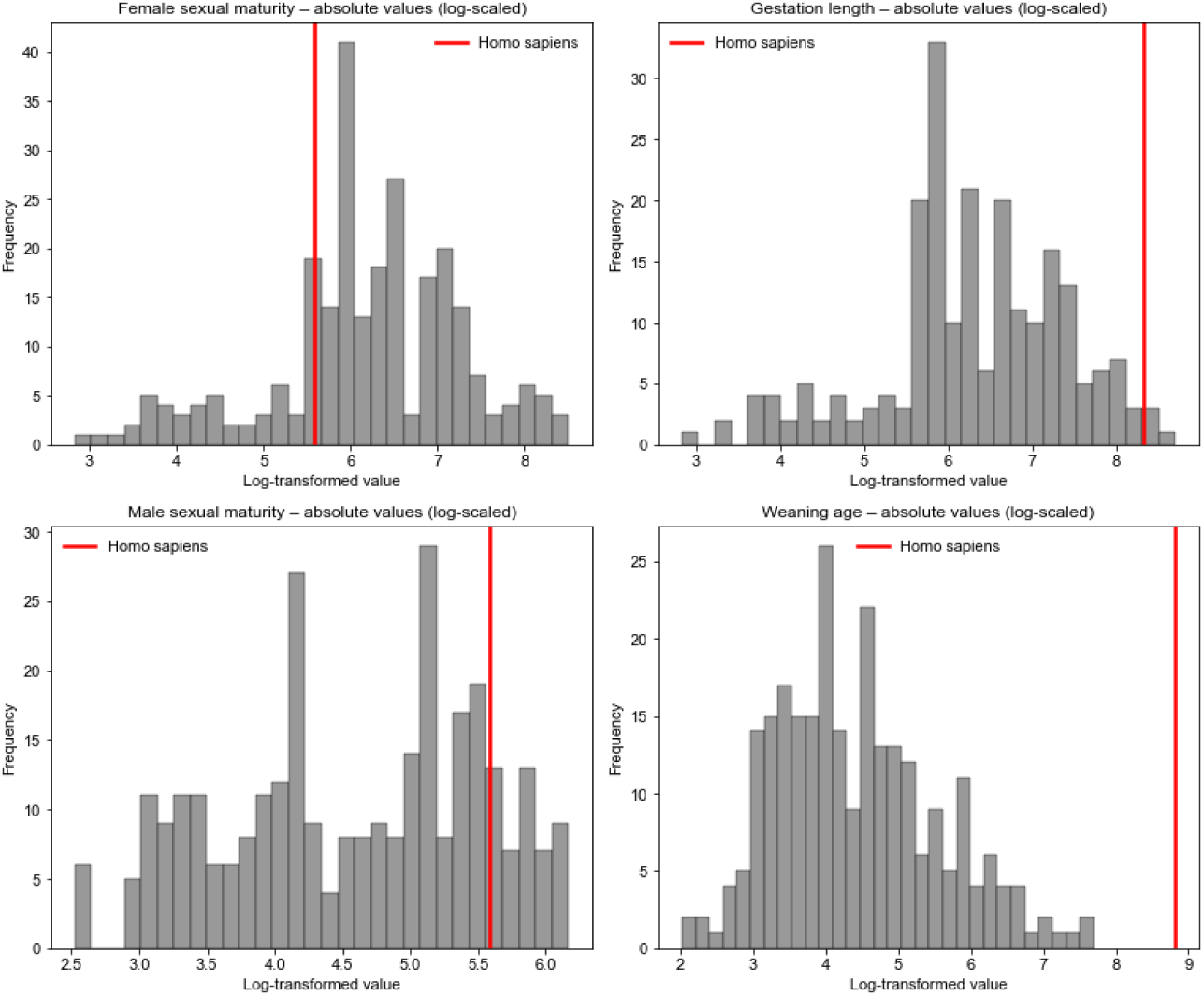
Distributions of absolute and proportional life history traits across the mammalian sample. The position of the human data point within each distribution is indicated by a vertical dashed line. Absolute values were log-transformed and proportional values were logit-transformed.

## 4. Discussion

Our study examined human developmental timing within a comparative framework, assessing gestation length, weaning age, and age at reproductive maturity as representatives of human dependency. We found that, as absolute metrics, all three life history traits showed humans to have extensive dependency periods across existing mammalian records, consistent with the view that human developmental timing is markedly protracted relative to most mammals. Humans are often characterized as unusually altricial, undergoing an exceptionally protracted period of postnatal dependency relative to other mammals (Portmann, 1969; Bogin, 1999), a feature tied to large brain size, secondary altriciality, and the cooperative provisioning thought to have shaped the evolution of hominin social organization and (Burkart et al., 2009; Isler & van Schaik, 2012).

Nevertheless, when expressed relative to the maximum lifespan of a given species, the pattern was less striking, showing low early-dependency measures and moderately high, but not exceptional reproductive maturation measures. This accords with recent arguments that several purportedly distinctive human life history features fall within the primate range once body size and phylogenetic position are controlled for (Robson et al., 2006; Miller et al., 2019). This might suggest that, although humans may still exhibit distinctive patterns along certain dimensions, such features appear more contingent in their emergence than previously assumed; extended dependency and related developmental timing events may not in themselves be sufficient to produce the cognitive and life history complexity observed in our species. Beyond primates, other long-lived mammals also exhibit slow life histories, although comparative investigations at this broader scale remain relatively limited. Asian and African elephants show late maturity and substantial post-reproductive lifespans (Lahdenperä et al., 2014; Lee et al., 2016), while several odontocete species such as killer whales, belugas, narwhals, and pilot whales exhibit menopause and decades of post-reproductive life (Foote, 2008; Ellis et al., 2018; Ellis et al., 2024). This suggests that, on the broader scheme, the evolution of developmental timing is shaped less by phylogeny than by a shared set of ecological and social conditions

Several limitations warrant consideration. The dataset was opportunistic and may bias results toward well-studied mammals, and the analyses are descriptive, taking no account of the phylogenetic relationships among species, which are substantial across mammalian life history traits (Blomberg et al., 2003).However, the principal aim here was to characterise the broad structure of mammalian developmental timing and to locate humans within it. Incorporating phylogenetic relatedness across a large group of mammalian lineages is nonetheless the natural next step and would permit firmer inferences about whether the convergences noted above are genuinely independent. Maximum captive lifespan, though useful for isolating intrinsic longevity from extrinsic mortality, may not reflect dynamics in natural settings, a caution that applies with particular force to humans, whose recorded maximum reflects modern conditions far removed from the higher-mortality settings under which human life history is thought to have evolved (Gurven & Kaplan, 2007). This qualifies the placement of humans on the proportional metrics rather than the broad ordering of taxa, which does not depend on the choice of longevity measure. Finally, the four traits examined here capture only a subset of the parameters relevant to offspring dependency. Adding metrics such as postnatal brain growth rate and neonatal metabolic requirements, alongside data from extant primates and the hominin fossil record, would meaningfully extend the present findings.

In conclusion, this study provides a broad comparative framework for evaluating human developmental timing across mammals. By assembling developmental data from a taxonomically diverse sample, we sought to establish a descriptive foundation against which claims of human life history exceptionalism can be evaluated more rigorously. Future studies integrating other metrics such as body size, phylogenetic comparative methods, and additional developmental traits will help determine which aspects of human dependency reflect general mammalian patterns and which represent derived features of the hominin lineage.

## Data Availability Statement

Data is available from the corresponding author upon reasonable request.

